# Hydrothermally treated yeast cell wall suppresses the growth of MCF7 cells under isolated conditions

**DOI:** 10.1101/2024.10.25.620192

**Authors:** Takanori Kitagawa

## Abstract

Traditionally, yeast cell wall (YCW) has limited applications because of their low solubility. To overcome this, a novel method was developed using a hydrothermal reaction to enhance its solubility and decrease its viscosity; this resulted in the production of a soluble form of YCW, known as the YCW treated with hydrothermal reaction (YCW-H), with broader chemical composition. Therefore, in the present study, we aim to investigate YCW-H, a hydrothermally treated YCW product with enhanced solubility, for the inhibition of cancer cell proliferation. YCW-H promotes plant growth by effectively regulating soil microspheres. YCW-H inhibited the growth of MCF-7 breast cancer cells even after complete physical separation from the cells. This suggests the presence of a diffusible cytotoxic factor in YCW-H, a phenomenon not observed in the presence of untreated YCW. Since YCW-H possess reactive carbon species (RCS), we hypothesized that the reactive radical species generated during the hydrothermal treatment of YCW are responsible for this effect. The addition of Fe(III) ions to YCW-H further amplified the production of RCS and its inhibitory activity across the plastic barrier, correlating with the increased RCS levels. The radicals migrated to the water in wells adjacent to YCW-H, indicating that RCS generation is a key determinant of the inhibition of cancer cell growth. Moreover, compounds containing RCS were confirmed to exhibit cytotoxicity. Thus, our findings underscore the RCS-containing compounds from YCW-H as promising candidates for developing novel medical devices for cancer treatment.

## Introduction

Beer is one of the most popular beverages globally, with an annual production of approximately 180 billion liters [1]. Fermentation with yeast, such as *Saccharomyces (S.) cerevisiae and S. pastorianus,* is a key step in beer brewing and generates a large amount of yeast cell residue [2-4]. This residue can be divided into two byproducts: yeast extract and cell wall, which are rich sources of proteins, minerals, vitamins, and polysaccharides, such as β-glucans [5]. Although yeast extract is used as a nutrient source, surplus amounts of the yeast cell wall (YCW) remain; the use of YCW is mostly limited to animal feed because it is difficult to process owing to its insolubility in water. To address this issue and make YCW a more useful material, we developed a new method using a hydrothermal reaction to increase its solubility and decreasing its viscosity [6]. The resulting soluble form of YCW is referred to as YCW treated with hydrothermal reaction (YCW-H). YCW-H exhibits enhanced solubility and potentially contains a broader spectrum of chemical components and properties than YCW. YCW-H is used as a plant fertilizer to control the bacterial biosphere by decreasing the oxidation–reduction potential in the soil [6]. Additionally, an enzyme-solubilized extract of YCW improves plant growth by inducing plant defense responses [7]. These findings indicate that the process of solubilizing YCW resulted in derivatives exhibiting distinct, previously unseen biological properties. Therefore, YCW-H serves as a water-soluble counterpart for YCW, enabling effective investigation of its biological effects and potential applications through *in vitro* assays.

YCW is emerging as a promising source of valuable polysaccharide components for various biomedical applications [8-11]. The major components of YCW include mannan oligosaccharides, β-glucans, and mannoproteins which are non-filamentous glycoproteins [12-14]. Various polysaccharides exhibit antitumor activity in addition to antioxidant, immunomodulatory, anti-inflammatory, and hypoglycemic activities [15-24]. The water-soluble polysaccharide fraction derived from mushroom mycelium exhibits antitumor effects by inhibiting cancer cell growth [25, 26]. Polysaccharides in YCW demonstrate a potential anticancer effect *in vivo* [22, 27]. In addition, YCW-H is used as a plant fertilizer, enhancing soil microsphere composition for optimal plant growth by converting unfavorable conditions into suitable ones [6]. This suggests that YCW-H modulates prokaryotic cell growth by selectively promoting or inhibiting the growth of certain bacterial species. Herein, we hypothesized that YCW-H could inhibit the proliferation of specific eukaryotic cells, such as cancer cells, through polysaccharides, polysaccharide derivatives, and/or chemicals solubilized via hydrothermal reactions with the aim to investigate a new application of YCW-H.

Here, we report that the YCW-H inhibited MCF7 cell growth, and this was observed even when YCW-H was placed outside the culture well. The growth inhibitory effect of YCW-H was transmitted via air and/or plastic materials, such as polystyrene (PS). Additionally, the growth inhibition via plastic positively correlated with the amount of reactive carbon species (RCS) in YCW-H, suggesting that a key factor governing the YCW-H-mediated inhibition of MCF7 cell growth is the generation of RCS.

## Results

### MCF7 cell growth was inhibited when cultured in a vessel adjacent to a well containing YCW-H

MCF7 breast cancer cells were treated with varying concentrations of YCW-H, ranging from 0–10%, to assess the growth-inhibitory effect of YCW-H. YCW-H inhibited MCF7 cell growth in a dose-dependent manner, with an EC50 of 1.8% (Fig. 1A), indicating potent anti-proliferative activity.

**Fig 1.**
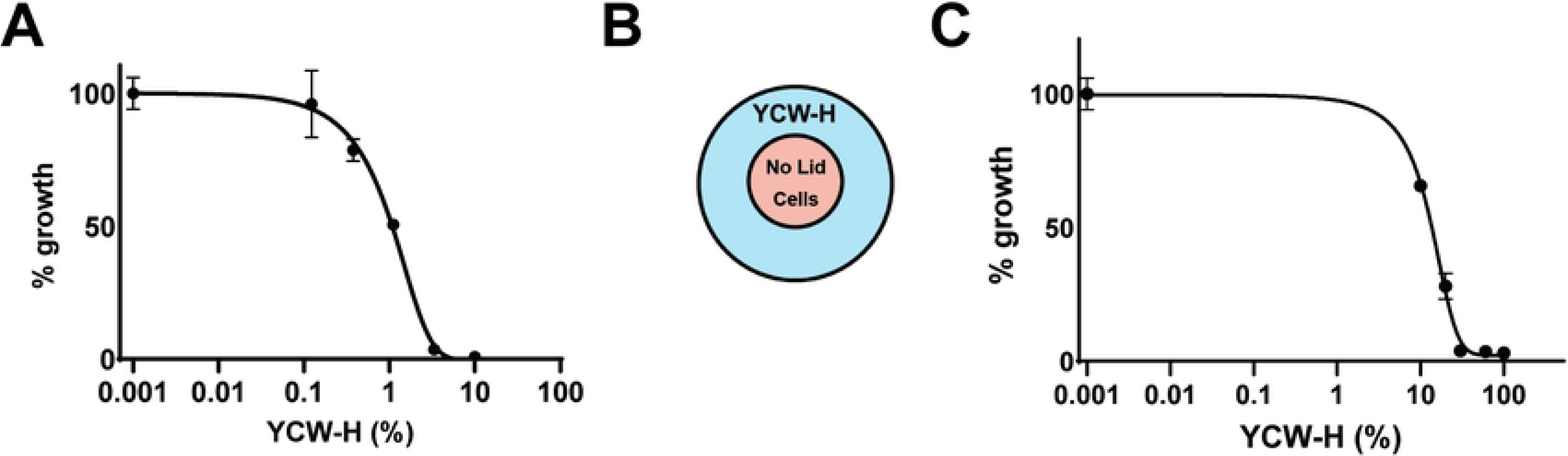
YCW-H suppressed cell growth from neighbor wells. All growth assays were performed with MCF7 cells. (**A)** Cell growth treated with various dose (0 and 10% to 0.123 % with 3 x dilution) of YCW-H was analyzed by curve fitting graph. Percentage (%) growth was calculated by the value of the wells without YCW-H as 100%. (**B)** Schematic diagram of the assay; MCF7 cells were seeded in 3.5 cm dish which located in 5 cm dish contained various concentrations of YCW-H. (**C)** MCF7 Cells with treatments of 0, 10, 20, 30, 60, and 100 % of YCW-H in its adjacent treatment without lid were analyzed cell growth by curve fitting graph. The growth assays here were performed by Crystal Violet assay. Percentage (%) growth was calculated by the value of PBS (0%) as 100%. Error bars are SD (n=3).

Cell growth assays revealed that wells lacking YCW-H but positioned adjacent to wells with high YCW-H concentrations exhibited reduced cell growth. To confirm this reduction, we performed growth assays using a two-layer culture system consisting of an inner dish placed in an outer dish (Fig. 1B). Varying concentrations of YCW-H were added to the outer dish. The cells were then cultured in the inner dish. A dose-dependent decrease in cell growth was observed, with an EC50 of 11.9% (Fig. 1C). These findings suggest that the airborne transmission of YCW-H inhibits MCF7 cell growth.

### Hydrothermal treatment conferred YCW with a growth-inhibitory effect on MCF7 cells

The growth inhibitory effects of YCW-H and YCW were compared to those of H₂O (control, 100%) to determine whether hydrothermal treatment was required. YCW was sterilized via autoclaving. MCF7 cells were seeded into three wells of a triple-well dish. After one day, the space around the culture wells was filled with H₂O, YCW-H, or YCW (Fig. 2A). Cell proliferation was significantly inhibited to 7.4% ± 0.32 in the presence of YCW-H compared to that of the control, while cells treated with YCW showed no significant inhibition (96.2% ± 8.1) (Fig. 2B). Therefore, the hydrothermal treatment of YCW endows it with the ability to inhibit cell growth.

**Fig 2.**
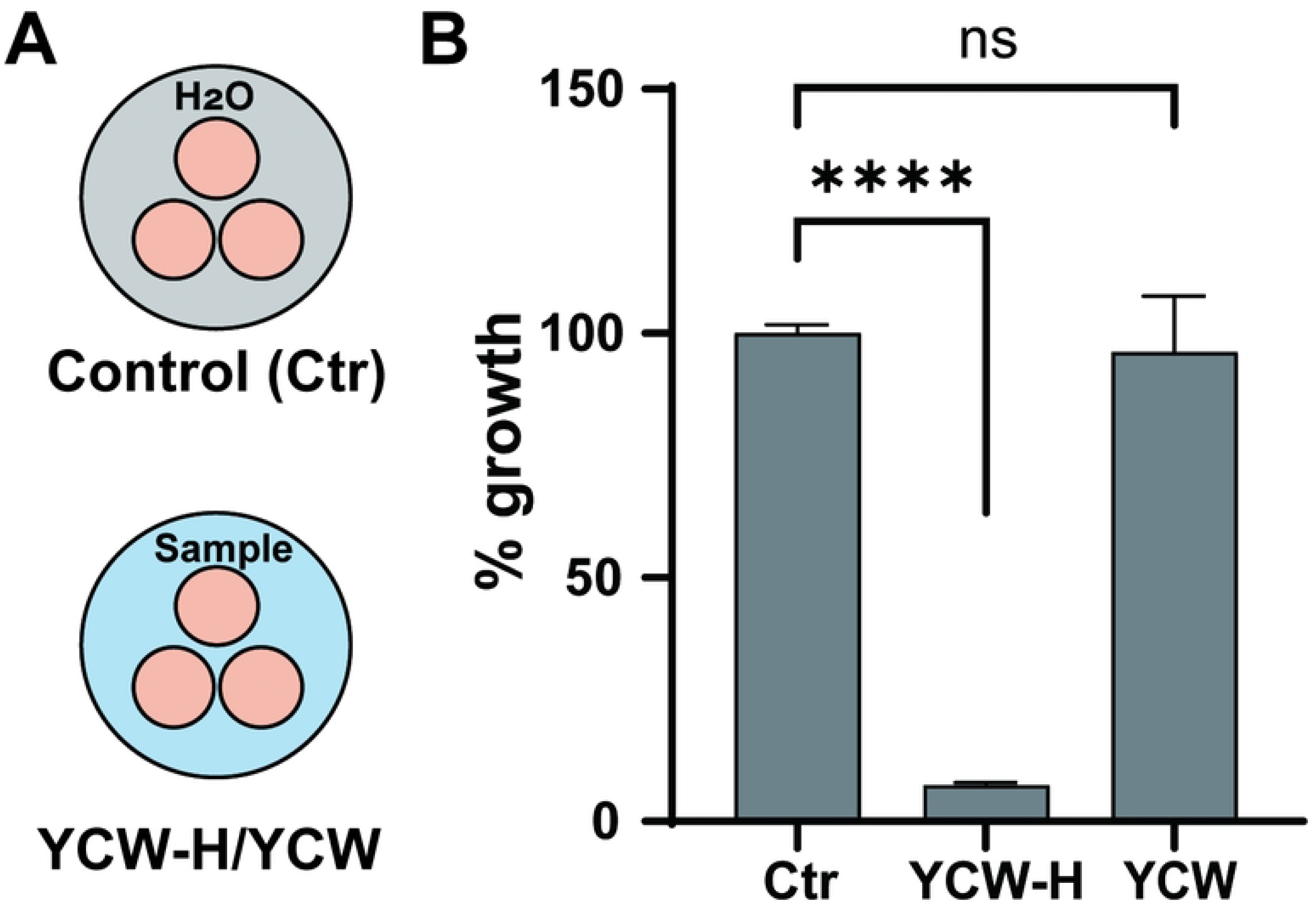
YCW-H acquired the ability of growth inhibition by treatment of hydrothermal reaction. **(A)** Schematic diagram of the assay; YCW-H/YCW or H_2_O (Control) were filled in a dish outside of triple wells. (**B)** Percentage (%) growth over a 3day culture was calculated by the value of control (Ctr) as 100%. Error bar is SD (n=3). p-value was indicated by ****: <0.0001 and ns: non-significance. Statistical significance was evaluated using a one-way ANOVA test as the result of the Brown-Forsythe test by Prism.

### YCW-H can inhibit MCF7 cell growth by trans-passing plastic walls

To further investigate whether the growth inhibition by YCW-H occurred via airborne transmission, a growth assay was performed under completely isolated conditions. MCF7 cells were seeded in a single-well dish, and the well was sealed with a coverslip using silicone grease to prevent air movement. The space outside the well was then filled with H₂O (control), YCW, or YCW-H (Fig. 3A, B). In this study, the adjacent wells that were treated were termed "closed treatment." Additionally, YCW-H significantly inhibited MCF7 cell growth to 58.7% ± 6.9 under closed treatment, compared with that observed for the control, while YCW had no effect on MCF7 cell growth (115.0% ± 10.8) (Fig. 3C). Thus, YCW-H inhibited cell growth even when the MCF7 cells were completely segregated from YCW-H.

**Fig 3.**
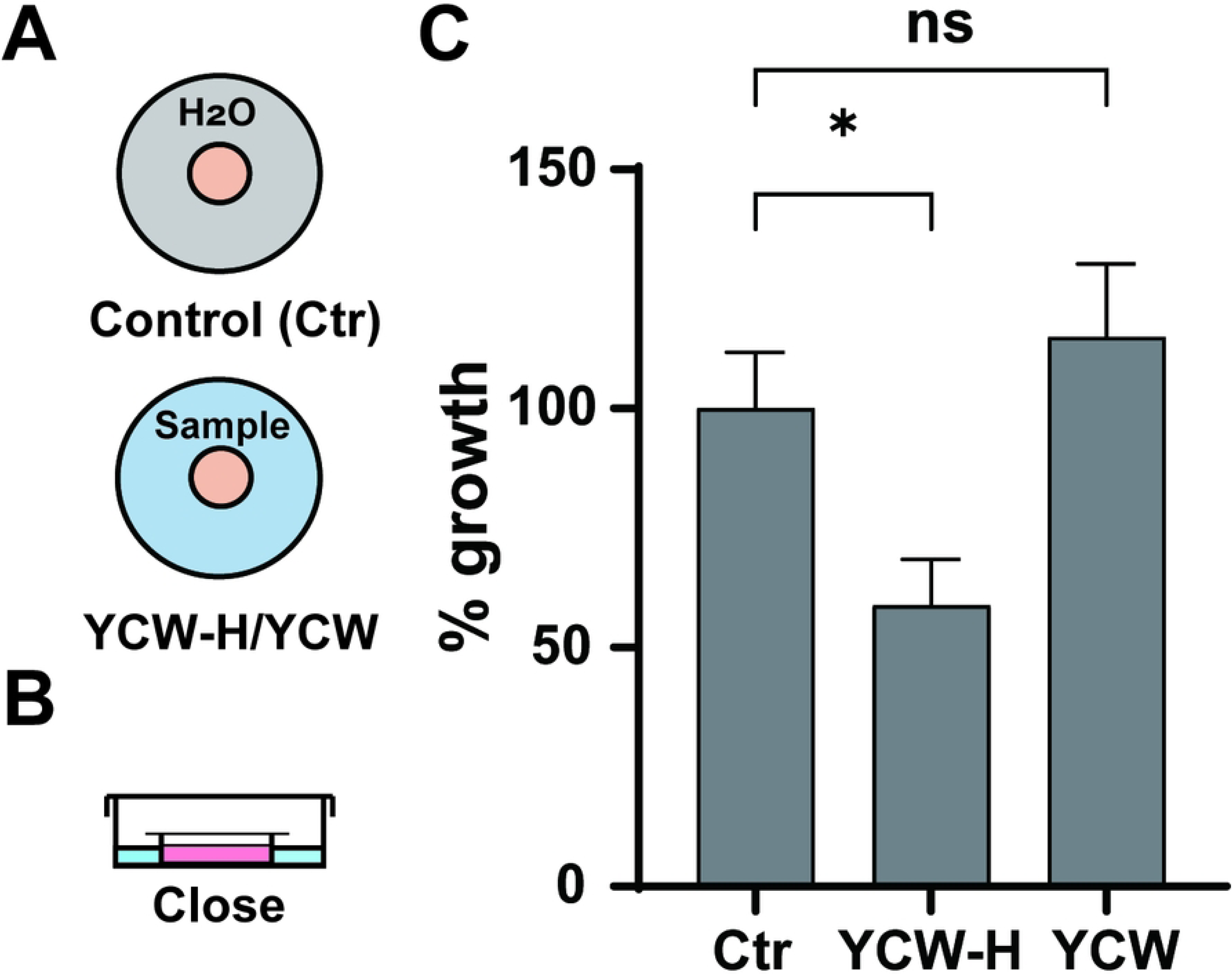
YCW-H suppressed cell growth from neighbor wells. **(A)** Schematic diagram of the assay; YCW-H/YCW or H_2_O (Control) were filled in a dish outside of a well. **(B)** The well was closed (Close) by the cover slide with silicon grease. (**C)** Percentage (%) growth for a 3 day culture was calculated by the value of control (Ctr) as 100%. Error bar is SD (n=3). p-value was indicated as *: P=0.016 and ns: non-significance: P=0.34. Statistical significance was evaluated using a one-way ANOVA test as the result of the Brown-Forsythe test by Prism.

### YCW-H contains reactive carbon species (RCS), the generation of which is regulated by iron ions

We hypothesize that YCW-H contains radical species that similarly influence cell proliferation, since studies show that reactive nitroxide species (RNS) prevent cell death through airborne transmission [28]. Therefore, electron spin resonance (ESR) spectroscopy was used to investigate the presence of radical species in YCW-H. The electronic g-factor, a key identifier for radical species, indicated that YCW-H contains RCS and Fe(III) complex, with g-factors of 2.003 and 4.25, respectively (Fig. 4A) [29-31]. The next step was to identify the inducer of RCS production in YCW-H. We investigated whether iron (Fe) ions could increase the RCS levels, since Fe ions can generate radicals, including RCS, in certain conditions [32-35]. The addition of FeSO_4_, Fe(II), to YCW-H increased the RCS levels to 2.7 μmol/L compared with 1.9 μmol/L that was observed in the presence of YCW-H alone (Fig. 4B, Table 1). Adding Fe_2_(SO_4_)_3_, “Fe(III),” into YCW-H increased the RCS level to 8.5 μmol/L, and a broad signal from hydrated Fe(III) was observed (Fig. 4C, Table 1). The concentration of Fe(III) complex increased from 0.15 to 0.29 mmol/L upon the addition of Fe(II), suggesting that Fe(II) oxidation increases Fe(III) levels. These findings suggest that Fe(II) and Fe(III) contribute to RCS production in the YCW-H.

**Fig 4.**
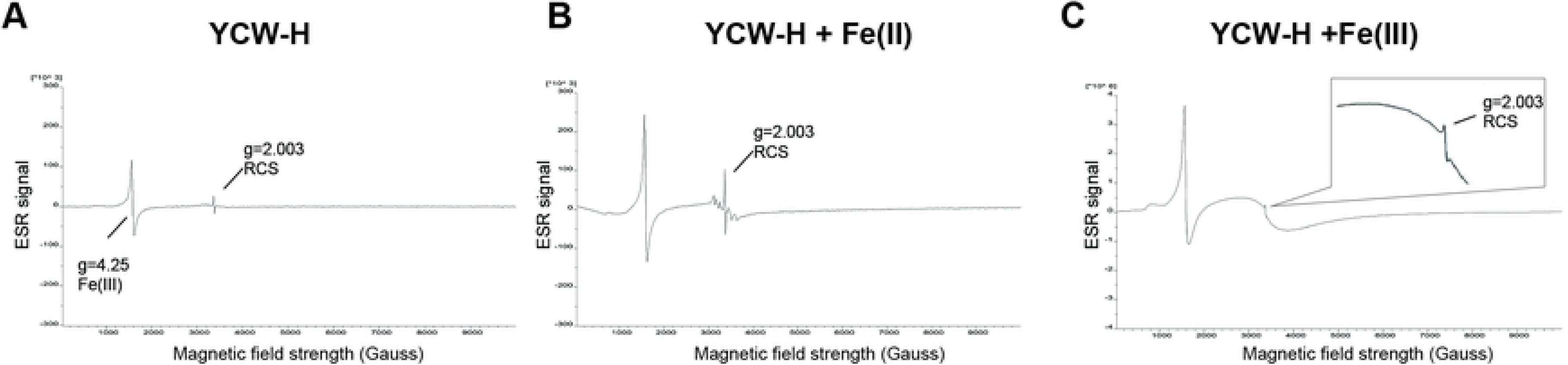
YCW-H possessed reactive carbon species. **(A)** YCW-H, **(B)** YCW-H and Fe (II) (46.3 mM) and (**C)** YCW-H and Fe (III) (46.3 mM) were analyzed by ESR. Magnified image of the RCS peak was shown in the box. X-axis indicates magnetic field strength (Gauss). Y-axis is units of the ESR signal. Value of a constant of proportionality, g, is the property of the electron.

**Table 1.** Concentration of RCS and Fe(III) complex measured by ESR analysis.

| Samples | Reactive carbon species: g=2.003 (µmol/L) | Fe(III) complex: g=4.25 (mmol/L) |
| --- | --- | --- |
| YCW-H | 1.9 | 0.15 |
| YCW-H + Fe(II) | 2.7 | 0.29 |
| YCW-H +Fe(III) | 8.5 | 11 |

The substantial change of about 10%, given the instrument's precision of ±5%, was deemed an effective difference.

### RCS can migrate through plastic walls during incubation

To investigate the migration of radical species to neighboring wells under closed conditions, ESR analysis was performed H₂O containing the spin trap N-tert-butyl-α-phenylnitrone immersed in YCW-H. ESR analysis revealed an increase in the concentration of radical adducts from 0.43 to 0.47 μmol/L. The addition of Fe₂(SO_4_)₃ (46.3 mM) to YCW-H further increased the adduct concentration to 0.51 μmol/L (Table 2). These findings suggest that radical species can permeate polystyrene walls, demonstrating their ability to migrate between completely separate compartments.

**Table 2.** Concentration of radical adducts with PBN measured by ESR analysis.

| Samples in tubes | Radical adducts (µmol/L) |
| --- | --- |
| H<sub>2</sub>O | 0.43 |
| YCW-H | 0.47 |
| YCW-H +Fe(III) | 0.51 |

The substantial change of about 10%, given the instrument's precision of ±5%, was deemed an effective difference.

### Ferrous and ferric ions accelerated MCF7 growth inhibition in YCW-H adjacent treatment under open and closed conditions

We hypothesized that the RCS in YCW-H might influence its inhibitory activity via plastic walls. Because Fe(II) and Fe(III) can increase the level of RCS, we investigated the effects of adding these ions on the growth-suppressive activity of YCW-H. Triple-well dishes were used for growth assays in the open treatment (Fig. 5A). YCW-H (25%) treatment alone inhibited growth to 48 ± 1.4% of the control; however, the addition of Fe(II) to YCW-H reduced the cell growth to 32 ± 1.1%, while the addition of Fe(III) reduced it to 24 ± 2.3%. This suggests that adding Fe(II) or Fe(III) to YCW-H accelerates the inhibition of cell proliferation compared to that of YCW-H alone under the open treatment conditions (Fig. 5A).

**Fig 5.**
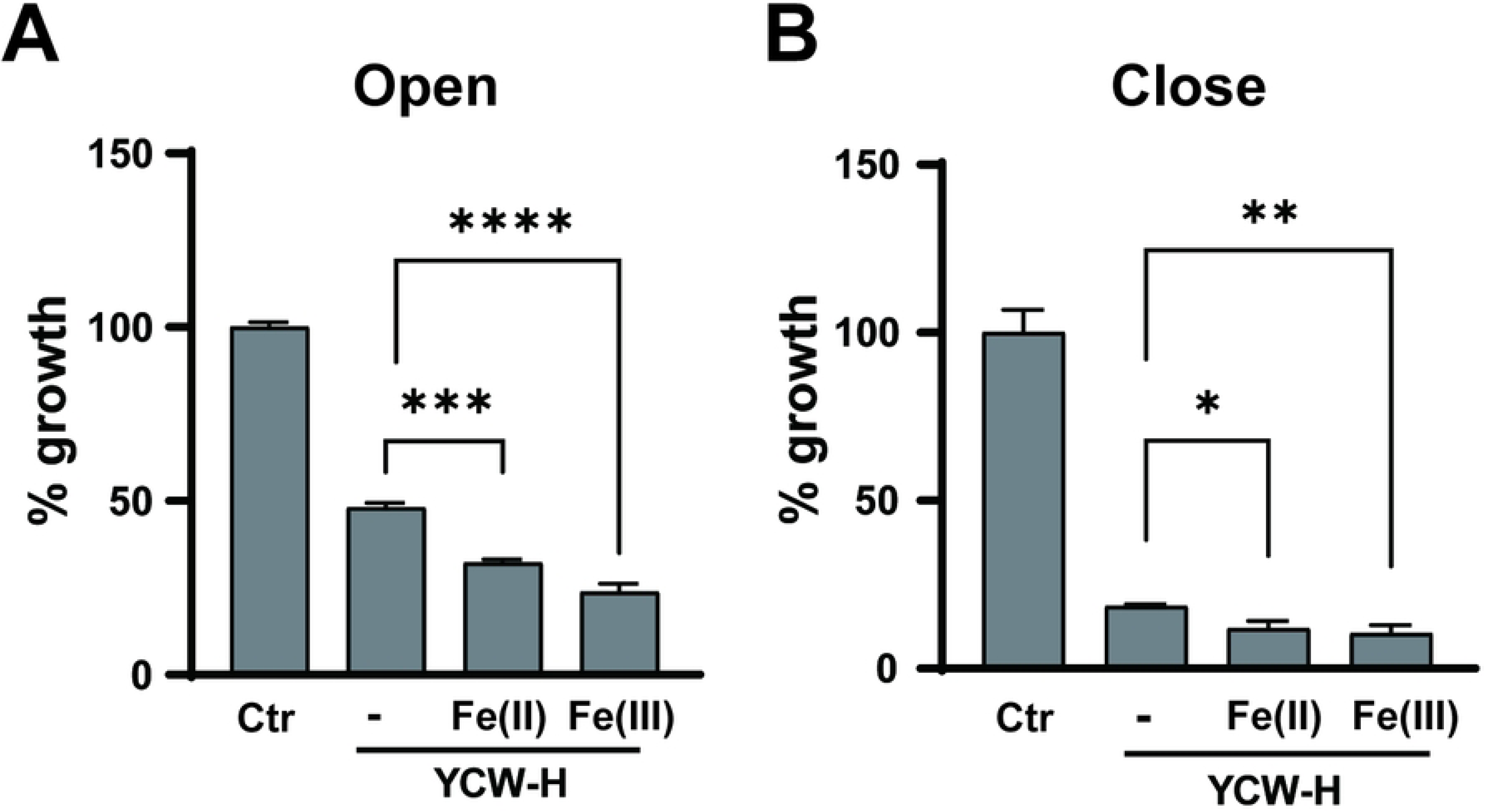
Fe(II) and Fe(III) were required to growth inhibition by YCW-H. **(A)** MCF7 cells were seeded in the wells of well dishes under open treatments. The outer space was filled with H_2_O (Ctr), and H_2_O (-), 40 mM of Fe(II) or Fe(III) in 25% YCW-H for 3 days. H_2_O alone was used as a control (100%). P-value is indicated as ****: P<0.0001, ***: P=0.0005. (**B)** MCF7 cells were seeded in the wells of well dishes under close treatments. The outer space was filled with H2O (Ctr), and H2O (-), 40 mM Fe(II), and 40 mM Fe(III) in YCW-H. Growth assays were performed by Crystal Violet assay. Percentage (%) growth was calculated by the value of H2O as 100%. Error bar is SD (n=3). P-value is indicated as**: P = 0.005 ** and *: P =0.01). Statistical significance was evaluated using a one-way ANOVA test as the result of the Brown-Forsythe test.

To further validate the effects of RCS via the combination of Fe(II) and Fe(III), the effects of the addition of Fe(II) and Fe(III) to YCW-H on MCF7 cell growth were explored. YCW-H alone inhibited proliferation by 18.6 ±0.46%, YCW-H with Fe(II) by 11.9 ±1.5%, and YCW-H with Fe(III) by 10.4 ± 1.8% compared with that of the control. This indicates that the addition of Fe(II) or Fe(III) enhances the growth-inhibitory ability of YCW-H against MCF7 cells under close treatment conditions (Fig. 5B). Additionally, the increase in RCS levels owing to the presence of Fe(II) and Fe(III) correlated with the extent of growth inhibition (Table 1). Thus, RCS activated by iron ions may play a key role in inhibiting MCF7 cell growth, even under conditions of complete separation.

### YCW-H was cytotoxic to MCF7 cells under open and closed YCW-H adjacent treatment conditions

To determine whether the growth suppression via YCW-H adjacent treatment was due to cytotoxicity, the percentage of live and dead cells in the treated cells was analyzed using flow cytometry with the detection reagent ViaCount^TM^. In open conditions, the presence of 25% YCW-H in the surrounding space resulted in 77.8 ± 2.3% of MCF7 cell death, compared to 6.8 ± 1.9% in the control group treated with H₂O (Fig. 6A). Under the close condition, 49.6 ± 3.1% of the treated cells underwent cell death, compared to 18.7 ± 3.0% of the control cells (Fig. 6B). Furthermore, at the exponential phase (Day 0), the percentage of dead cells was 18 ± 1.5%, similar to the percentage of untreated cells, suggesting that the closed treatment did not affect MCF7 cell proliferation (Fig. 6B). Reactive oxygen species (ROS) levels in the YCW-H adjacent treatment were examined under closed treatment conditions, since ROS activation induces cell death [34]. ROS were elevated in cells subjected to YCW-H closed treatment before cell death induction but not in untreated cells (Fig. 6C). The increase in ROS levels correlated with the number of dead cells, suggesting that ROS activation could be involved in RCS-mediated cell death. Therefore, the growth inhibition observed in the YCW-H adjacent treatments was deemed to be the result of cytotoxicity.

**Fig. 6.**
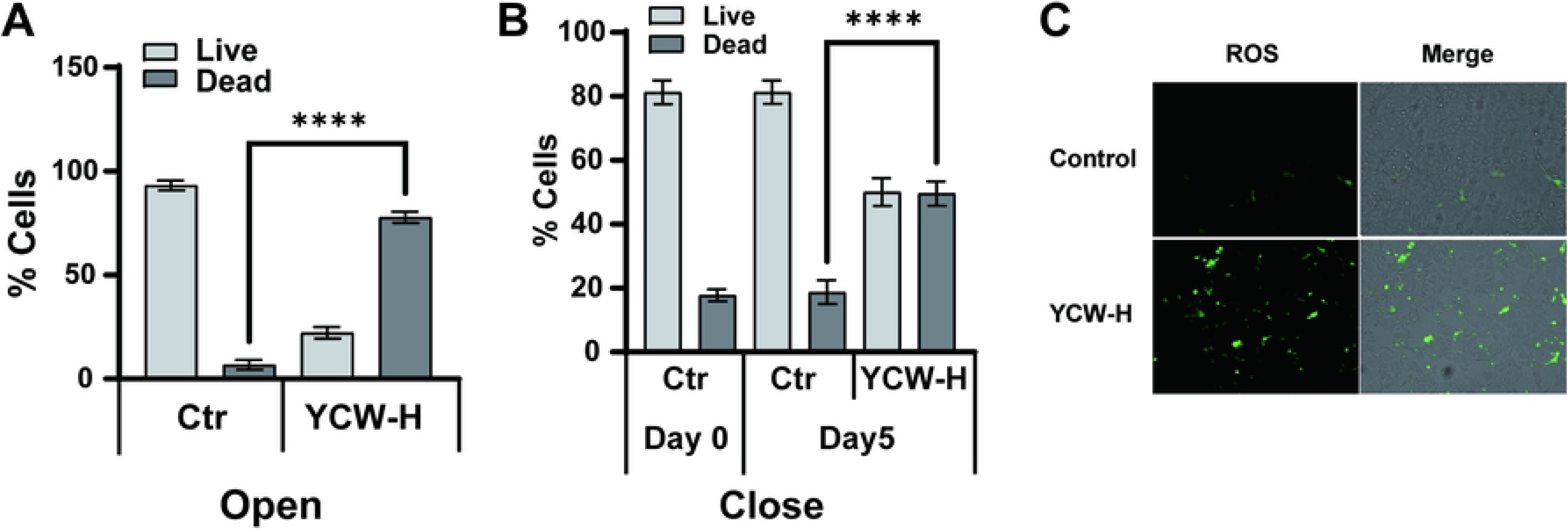
YCW-H adjacent treatment induced cell death and ROS. **(A)** Number of live and dead cells with and without YCW-H were measured by Guava® with ViaCount™. The cells were cultured for 4 days under Open condition. Error bar is SD (n=3). P-value is indicated as ****: p<0.0001. (**B)** Number of live and dead cells with and without YCW-H after 5 days in Close condition. Day0 is one day before the treatment was started. Error bar is SD (n=3). P-value is indicated as ****; p<0.0001. Statistical significance was evaluated using a one-way ANOVA test as the result of the Brown-Forsythe test. (**C)** ROS induction was detected by ROS detection kit. MCF7 cells were cultured with and without YCW-H treatment for 3 days in close condition. Green fluorescence indicated ROS induction in cells (ROS). Images of brightfield and fluorescence were overlaid (Overlay). Representative Images from triplicate assays were indicated.

## Discussion

Our study investigated the effects of YCW-H on the growth of cancer cells. The results indicated that YCW-H induces cytotoxicity in cancer cells regardless of their physical proximity, such as completely isolated culture wells. RCS generated by YCW-H is a key contributor to its growth-inhibitory effects via a plastic wall; however, the specific nature of these radicals remains unknown. Research on the biological importance of RCS-containing compounds in regulating cell proliferation remains limited [36-38]. However, carbon-centered radicals are involved in the induction of apoptotic cell death [39]. In addition, alkyl radicals, one of RCS, are implicated in ferroptosis and possess cytotoxicity to tumor cells [38]. Thus, YCW-H exhibits unique properties that certain chemical substances with RCS migrate through the plastic wall.

ESR analysis revealed that YCW-H contains ferric ions as Fe(III) complexes. USDA data shows that barley malt has 4.7 mg of iron per 100 g, suggesting that barley malts is a potential iron-source in YCW-H. Furthermore, *S. cerevisiae* uses mannoproteins present in the cell wall to facilitate the uptake of Fe(III) complexes, thereby accumulating more Fe(III) in the cell wall [40, 41]. Thus, the cell walls of the yeast may retain a sufficient amount of Fe(III) and organic carbon after fermentation, forming Fe(III) complexes.

Our result indicated that YCW-H possessed RCS. The presence of Fe(III) enhanced the RCS-generation activity of YCW-H; the induction of RCS generation proportionally enhanced the growth-inhibitory effects of YCW-H under open and closed conditions. Adding Fe(II) significantly promotes growth inhibitory effect of MCF7 cells, as well as increases the amount of Fe(III) complexes and RCS. The effects of Fe (II) addition may be attributed to the elevated formation of Fe(III). This elevation of Fe(III) results from a redox reaction between Fe(II) and Fe(III), which can be exploited for growth inhibition. Fe(II) suspensions, when exposed to O_2_ under certain conditions, undergo spontaneous oxidation at room temperature [42]. In contrast, the reduction of Fe(III) to Fe(II) occurs with extremely low efficiency owing to its minimal redox potential [42]. Thus, Fe(II) supplementation in YCW-H catalyzes the conversion to Fe(III), enhancing YCW-H-dependent cell death via inducing RCS. These findings support the unique growth inhibitory effect of Fe(III)-associated RCS and suggest that Fe(III) enhances the activity of YCW-H.

This study shows that the inhibition of cell growth by YCW-H positively correlates with an increase in RCS levels. A YCW-H treatment in adjacent wells increases the intracellular ROS levels, leading to cell death, while the factors responsible for activating ROS and cell death remain unknown. Cellular ROS are endogenously produced during mitochondrial oxidative phosphorylation in response to various biological processes, such as xenobiotic metabolism, phagocytosis, and arginine metabolism [43-45]. Excessive cellular ROS can damage key cellular components, including proteins, nucleic acids, and organelles, which may ultimately trigger cell death [46]. Both intrinsic and extrinsic pathways converge on ROS-mediated damage, making ROS a critical factor in cancer cell death [47]. Peroxyl radicals (ROO•), formed by the direct reaction of oxygen with alkyl radicals (R•), are implicated in ferroptosis via lipid ROS controlled by singlet oxygen, supporting our finding [38]. Therefore, RCS-containing compounds in the YCW-H adjacent treatments could induce intracellular ROS production, leading to cell death.

Our findings indicate that the RCS-containing factors of YCW-H may migrate through the plastic walls of polystyrene culture plates. In this case, chemical transport in the polystyrene wall would be necessary. The migration requires three steps: absorption into the polystyrene, diffusion through the material, and finally, release into the adjacent well. Chemicals are well known to be absorbed into polystyrene assay plates, since this chemical adsorption to polystyrene is often responsible for most of the reduction in the levels of the observed chemicals in the medium in *in vitro* assays by uneven influx and efflux rates within the same well [48-52]. For the transmigration to a new well, the chemical diffusion in polystyrene would occur through chemical gradient between the two wells driven by chemical concentrations in each media. The chemicals would be eventually released to a fresh medium, when the chemicals reached to the side facing it. Alternatively, RCS would be transferred by radical chain reaction, since thermal degradation of most of the polymers is a typical radical chain mechanism with RCS [53]. Therefore, the chemicals with RCS-containing factors could be transferred across the plastic wall.

In conclusion, the RCS from YCW-H, enhanced by iron ions, exhibited potent anti-proliferative and pro-apoptotic effects, suggesting its potential as a therapeutic agent for cancer. Chemotherapy, commonly used in the treatment of various cancers, including skin and breast cancer, often leads to side effects such as hair loss, diarrhea, vomiting, chest pain, constipation, difficulty breathing, fatigue, mucositis, and rash. To improve the patient’s quality of life, future investigation should focus on developing innovative therapies that minimize these side effects. The agents capable of overcoming plastic walls represent a novel approach to cell inhibition and hold potential for the development of noninvasive medical devices for cancer treatment.

## Materials and Methods

### Cell culture

The breast cancer cell lines, MCF7, were purchased from the Japanese Collection of Research Bioresources Cell Bank (JCRB, Osaka Japan). MCF7 cells were cultured in DMEM with 10% fetal bovine serum and 1% penicillin and streptomycin at 37°C in 5% CO2. Regular testing of mycoplasma contamination was performed in the cell lines using CycleavePCR™ Mycoplasma Detection Kit (Takara, Shiga Japan), and only mycoplasma free cells were used for experiments.

### Preparation of YCW

Yeast cell walls (ASAHI Group Foods Ltd., Tokyo Japan) are made into a 15% (w/v) solution with water and then sterilized by autoclave at 121 °C for 20 min.

### Hydrothermal reaction of YCW - YCW-H

Yeast cell walls (ASAHI Group Foods Ltd., Tokyo Japan) are made into a 15% (w/v) solution with water. The solution was gradually heated in an industrial autoclave from room temperature to a temperature of 180 °C and pressure of 1.6 MPa over 2 hours. Once the temperature and pressure reached that level, stopped the autoclave immediately and allowed the solution to return to room temperature naturally.

### Cell growth assay

Cell growth was measured by crystal violet assay. Cells were fixed with 100 μL of 1% paraformaldehyde #163-20145 (WAKO, Osaka Japan), and stained with 100 μL of 0.2% crystal violet # 15192 (MUTO PURE CHEMICALS, Tokyo Japan) after cell cultures were completed. The crystal violet was extracted by 100 μL of 50% ethanol. The absorption of the extracts was measured at 570 nm using a SpectraMax® M3 (Molecular Devices, San Jose CA USA). *Cell culture for 96 well growth assay*; MCF7 cell lines were seeded 4,000 cells in each well of 96 well plate and incubate for one days in 100 μL of DMEM supplemented with 10%FBS, 1% penicillin and streptomycin at 37°C in 5% CO2. 100 μL of DMEM with 10%FBS, 1% penicillin and streptomycin with and without YCW-H were added in the cell culture. Various percentages of YCW-H were diluted in DMEM. The cells were subsequently incubated for 3-5 days. *Cell culture with adjacent treatment under open condition in culture dishes*; 8×10^4^ MCF7 cells were seeded in 3.5 cm dish which located in 5 cm dish contained various concentrations of YCW-H. Varying concentrations of YCW-H which were diluted in PBS — 10, 20, 30, 60, and 100%—as well as PBS (0%), were added to the outer dish. *Cell culture with adjacent treatment under open and close condition in well plates;* MCF7 cell lines were seeded 4,000 cells in each well (*ϕ*8 mm) on a triple well dish (*ϕ*35 mm) # 3970-103 (AGC/Iwaki, Shizuoka Japan) and incubated for one days in 100 μL of DMEM supplemented with 2%FBS, 1% penicillin and streptomycin at 37°C in 5% CO2. For the open culture condition, the cells were subsequently incubated with and without 2 mL of YCW-H (25%) in the outside space of triple-wells for 3-5 days. For the YCW-H adjacent treatment, “close” condition, the wells, in which cells are cultured, were sealed by silicone grease with *ϕ*13 mm cover slides and then incubated with and without YCW-H in the outside space of a well for 3-5 days.

### Cell counts for live and dead cells

To harvest adherent cells, cells were detached with trypsin and mixed with the culture medium. All the cell mixtures were resuspended at 1 x 10^6^ cells/mL with culture medium. 380 μL of ViaCount reagent # SKU 4000-0040 (Cytek Biosciences, Fremont CA USA) were added to 20 μL of the cell suspension and incubated at 25°C for 10 minutes. Cell counting was performed automatically by Guava® easyCyte^TM^ 5HT and the analyses were provided plots separating live and dead cells by the Guava® ViaCount™ Software Module (Merck Millipore, Burlington MA USA).

### ROS detection

ROS Assay Kit -Highly Sensitive DCFH-DA # R252 (Dojindo, Kumamoto Japan) was used to detect intercellular ROS activation. MCF7 cells (5,000 cells) were seeded into triple well dish # 3970-103 (AGC/Iwaki, Shizuoka Japan) and incubated with and without YCW-H in the outside space of triple-wells for 3 days under close conditions. After the culture medium was removed, cells were stained with DCFH-DA Dye following the manual. Fluorescence signals were imaged by 20x lens with BZ-X710 fluorescence microscope (Keyence, Osaka Japan).

### Rapid-Freezing Method for ESR analysis

YCW-H (600 μL) was rapidly frozen by squirting it through a needle (i.d. 0.7mm) into the cooling liquid at about 20 K according to the method reported previously [54]. A series of cocktails of YCW-H mixed with H2O, Fe(SO4) (46.3 mM), or Fe2(SO4)3 (46.3 mM) were used for analysis.

### ESR measurements

The samples were analyzed by EMXplus ESR spectrometer (Bruker, Billerica, MA USA) using an X-band standard frequency of 8.8–9.6 GHz. To identify the peaks, the signal components were analyzed by the ES-IPRITS data system with version 3.00 analysis software installed in the ESR instrument. The following ESR parameters were used: a frequency of 9.42 GHz, center field of 335 ± 10 mT, modulation frequency of 100 kHz, time constant of 0.03 s, and power of 5.00 mW. Aqueous samples were loaded into a LC-12 aqueous quartz flat cell (JEOL)17.

### Preparation of H₂O containing the spin trap N-tert-butyl-α-phenylnitrone immersed in YCW-H

YCW-H was placed in a 15-mL polystyrene tube and immersed 5-mL polystyrene tube containing 500 μL of H₂O with 2 μM of the spin trap N-tert-butyl-α-phenylnitrone for 3 days.

## Acknowledgements

I am grateful to Director Kazuyuki Otsuka of the Osaka Bio-medical College (OBM) Research Center for his invaluable technical advice and insightful suggestions. I would also like to express my sincere thanks to Drs. Koichi Okumura and Tetsuya Moriyama of the OBM Research Center for their enlightening discussions, which significantly contributed to the improvement of this manuscript.

## References

1. 1. Barth SJ. BarthHaas Report. 2022.

2. Dequin S, Casaregola S. The genomes of fermentative Saccharomyces. C R Biol. 2011;334(8-9):687–93. Epub 20110701. doi: 10.1016/j.crvi.2011.05.019. PubMed PMID: 21819951.

3. Rainieri S, Kodama Y, Kaneko Y, Mikata K, Nakao Y, Ashikari T. Pure and mixed genetic lines of Saccharomyces bayanus and Saccharomyces pastorianus and their contribution to the lager brewing strain genome. Appl Environ Microbiol. 2006;72(6):3968–74. doi: 10.1128/AEM.02769-05. PubMed PMID: 16751504; PubMed Central PMCID: PMCPMC1489639.

4. Tamai Y, Momma T, Yoshimoto H, Kaneko Y. Co-existence of two types of chromosome in the bottom fermenting yeast, Saccharomyces pastorianus. Yeast. 1998;14(10):923–33. doi: 10.1002/(SICI)1097-0061(199807)14:10<923::AID-YEA298>3.0.CO;2-I. PubMed PMID: 9717238.

5. Marson GV, de Castro RJS, Belleville MP, Hubinger MD. Spent brewer’s yeast as a source of high added value molecules: a systematic review on its characteristics, processing and potential applications. World J Microbiol Biotechnol. 2020;36(7):95. Epub 20200624. doi: 10.1007/s11274-020-02866-7. PubMed PMID: 32583032.

6. Kitagawa T. Seibutu-kougaku Kaishi (Japanese). 2018;96:461.

7. Narusaka M, Minami T, Iwabuchi C, Hamasaki T, Takasaki S, Kawamura K, et al. Yeast cell wall extract induces disease resistance against bacterial and fungal pathogens in Arabidopsis thaliana and Brassica crop. PLoS One. 2015;10(1):e0115864. Epub 20150107. doi: 10.1371/journal.pone.0115864. PubMed PMID: 25565273; PubMed Central PMCID: PMCPMC4286235.

8. Bastos R, Oliveira PG, Gaspar VM, Mano JF, Coimbra MA, Coelho E. Brewer’s yeast polysaccharides - A review of their exquisite structural features and biomedical applications. Carbohydr Polym. 2022;277:118826. Epub 20211028. doi: 10.1016/j.carbpol.2021.118826. PubMed PMID: 34893243.

9. Liu Y, Wu Q, Wu X, Algharib SA, Gong F, Hu J, et al. Structure, preparation, modification, and bioactivities of beta-glucan and mannan from yeast cell wall: A review. Int J Biol Macromol. 2021;173:445–56. Epub 20210123. doi: 10.1016/j.ijbiomac.2021.01.125. PubMed PMID: 33497691.

10. Ferreira IMPLVO, Pinho O, Vieira E, Tavarela JG. Brewer’s Saccharomyces yeast biomass: characteristics and potential applications. Trends in Food Science & Technology. 2010;21(2):77–84. doi: 10.1016/j.tifs.2009.10.008.

11. Puligundla P, Mok C, Park S. Advances in the valorization of spent brewer’s yeast. Innovative Food Science & Emerging Technologies. 2020;62:102350. doi: 10.1016/j.ifset.2020.102350.

12. Cabib E, Roberts R, Bowers B. Synthesis of the yeast cell wall and its regulation. Annu Rev Biochem. 1982;51:763–93. doi: 10.1146/annurev.bi.51.070182.003555. PubMed PMID: 7051965.

13. Cawley TN, Ballou CE. Identification of two Saccharomyces cerevisiae cell wall mannan chemotypes. J Bacteriol. 1972;111(3):690–5. doi: 10.1128/jb.111.3.690-695.1972. PubMed PMID: 4559821; PubMed Central PMCID: PMCPMC251341.

14. Kollar R, Reinhold BB, Petrakova E, Yeh HJ, Ashwell G, Drgonova J, et al. Architecture of the yeast cell wall. Beta(1-->6)-glucan interconnects mannoprotein, beta(1-->)3-glucan, and chitin. J Biol Chem. 1997;272(28):17762-75. doi: 10.1074/jbc.272.28.17762. PubMed PMID: 9211929.

15. Chaisuwan W, Phimolsiripol Y, Chaiyaso T, Techapun C, Leksawasdi N, Jantanasakulwong K, et al. The Antiviral Activity of Bacterial, Fungal, and Algal Polysaccharides as Bioactive Ingredients: Potential Uses for Enhancing Immune Systems and Preventing Viruses. Front Nutr. 2021;8:772033. Epub 20211105. doi: 10.3389/fnut.2021.772033. PubMed PMID: 34805253; PubMed Central PMCID: PMCPMC8602887.

16. Guo R, Chen M, Ding Y, Yang P, Wang M, Zhang H, et al. Polysaccharides as Potential Anti-tumor Biomacromolecules -A Review. Front Nutr. 2022;9:838179. Epub 20220228. doi: 10.3389/fnut.2022.838179. PubMed PMID: 35295918; PubMed Central PMCID: PMCPMC8919066.

17. Jiang Y, Zhou W, Zhang X, Wang Y, Yang D, Li S. Protective Effect of Blood Cora Polysaccharides on H9c2 Rat Heart Cells Injury Induced by Oxidative Stress by Activating Nrf2/HO-1 Signal Pathway. Front Nutr. 2021;8:632161. Epub 20210302. doi: 10.3389/fnut.2021.632161. PubMed PMID: 33738296; PubMed Central PMCID: PMCPMC7960668.

18. Muszynska B, Grzywacz-Kisielewska A, Kala K, Gdula-Argasinska J. Anti-inflammatory properties of edible mushrooms: A review. Food Chem. 2018;243:373–81. Epub 20170930. doi: 10.1016/j.foodchem.2017.09.149. PubMed PMID: 29146352.

19. Nauts HC, Swift WE, Coley BL. The treatment of malignant tumors by bacterial toxins as developed by the late William B. Coley, M.D., reviewed in the light of modern research. Cancer Res. 1946;6:205–16. PubMed PMID: 21018724.

20. Pillemer L, Ross OA. Alterations in serum properdin levels following injection of Zymosan. Science. 1955;121(3151):732-3. doi: 10.1126/science.121.3151.732. PubMed PMID: 14372981.

21. Ren L, Zhang J, Zhang T. Immunomodulatory activities of polysaccharides from Ganoderma on immune effector cells. Food Chem. 2021;340:127933. Epub 20200826. doi: 10.1016/j.foodchem.2020.127933. PubMed PMID: 32882476.

22. Suzuki I, Itani T, Ohno N, Oikawa S, Sato K, Miyazaki T, et al. Effect of a polysaccharide fraction from Grifola frondosa on immune response in mice. J Pharmacobiodyn. 1985;8(3):217–26. doi: 10.1248/bpb1978.8.217. PubMed PMID: 3891963.

23. Zhao R, Gao X, Cai Y, Shao X, Jia G, Huang Y, et al. Antitumor activity of Portulaca oleracea L. polysaccharides against cervical carcinoma in vitro and in vivo. Carbohydr Polym. 2013;96(2):376–83. Epub 20130417. doi: 10.1016/j.carbpol.2013.04.023. PubMed PMID: 23768576.

24. Zhou WJ, Wang S, Hu Z, Zhou ZY, Song CJ. Angelica sinensis polysaccharides promotes apoptosis in human breast cancer cells via CREB-regulated caspase-3 activation. Biochem Biophys Res Commun. 2015;467(3):562–9. Epub 20150930. doi: 10.1016/j.bbrc.2015.09.145. PubMed PMID: 26431878.

25. Cui FJ, Tao WY, Xu ZH, Guo WJ, Xu HY, Ao ZH, et al. Structural analysis of anti-tumor heteropolysaccharide GFPS1b from the cultured mycelia of Grifola frondosa GF9801. Bioresour Technol. 2007;98(2):395–401. Epub 20060203. doi: 10.1016/j.biortech.2005.12.015. PubMed PMID: 16459075.

26. Do TTH, Lai TNB, Stephenson SL, Tran HTM. Cytotoxicity activities and chemical characteristics of exopolysaccharides and intracellular polysaccharides of Physarum polycephalum microplasmodia. BMC Biotechnol. 2021;21(1):28. Epub 20210327. doi: 10.1186/s12896-021-00688-5. PubMed PMID: 33773573; PubMed Central PMCID: PMCPMC8005236.

27. Fortin O, Aguilar-Uscanga B, Vu KD, Salmieri S, Lacroix M. Cancer Chemopreventive, Antiproliferative, and Superoxide Anion Scavenging Properties of Kluyveromyces marxianus and Saccharomyces cerevisiae var. boulardii Cell Wall Components. Nutr Cancer. 2018;70(1):83–96. Epub 20171116. doi: 10.1080/01635581.2018.1380204. PubMed PMID: 29144773.

28. Mizuno H, Kubota C, Takigawa Y, Shintoku R, Kannari N, Muraoka T, et al. 2,2,6,6-Tetramethylpiperidine-1-oxyl acts as a volatile inhibitor of ferroptosis and neurological injury. The Journal of Biochemistry. 2022;172(2):71-8. doi: 10.1093/jb/mvac044.

29. Hirota Y, Haida M, Mohtarami F, Takeda K, Iwamoto T, Shioya S, et al. Implication of ESR signals from ceruloplasmin (Cu(2+)) and transferrin (Fe(3+)) in pleural effusion of lung diseases. Pathophysiology. 2000;7(1):41–5. doi: 10.1016/s0928-4680(99)00033-4. PubMed PMID: 10825684.

30. Kono Y, Kashine S, Yoneyama T, Sakamoto Y, Matsui Y, Shibata H. Iron chelation by chlorogenic acid as a natural antioxidant. Biosci Biotechnol Biochem. 1998;62(1):22–7. doi: 10.1271/bbb.62.22. PubMed PMID: 9501514.

31. Tian L, Koshland CP, Yano J, Yachandra VK, Yu IT, Lee SC, et al. Carbon-Centered Free Radicals in Particulate Matter Emissions from Wood and Coal Combustion. Energy Fuels. 2009;23(5):2523–6. Epub 20090327. doi: 10.1021/ef8010096. PubMed PMID: 19551161; PubMed Central PMCID: PMCPMC2700017.

32. Arai N, Narasaka K. Development of New Methods for Generation of Radical Species by One-Electron Oxidation with Metallic Oxidants toward Construction of Carbon Skeletons. Journal of Synthetic Organic Chemistry, Japan. 1996;54(11):964–75. doi: 10.5059/yukigoseikyokaishi.54.964.

33. Jiang H, Lai W, Chen H. Generation of Carbon Radical from Iron-Hydride/Alkene: Exchange-Enhanced Reactivity Selects the Reactive Spin State. ACS Catalysis. 2019;9(7):6080–6. doi: 10.1021/acscatal.9b01691.

34. Klebanoff SJ, Waltersdorph AM, Michel BR, Rosen H. Oxygen-based free radical generation by ferrous ions and deferoxamine. J Biol Chem. 1989;264(33):19765–71. PubMed PMID: 2555330.

35. Feng Y, Wu D, Li H, Bai J, Hu Y, Liao C, et al. Activation of Persulfates Using Siderite as a Source of Ferrous Ions: Sulfate Radical Production, Stoichiometric Efficiency, and Implications. ACS Sustainable Chemistry & Engineering. 2018;6(3):3624–31. doi: 10.1021/acssuschemeng.7b03948.

36. Ning S, Liu Z, Chen M, Zhu D, Huang Q. Nanozyme hydrogel for enhanced alkyl radical generation and potent antitumor therapy. Nanoscale Adv. 2022;4(18):3950–6. Epub 20220816. doi: 10.1039/d2na00395c. PubMed PMID: 36133353; PubMed Central PMCID: PMCPMC9470029.

37. Seren S, Joly JP, Voisin P, Bouchaud V, Audran G, Marque SRA, et al. Neutrophil Elastase-Activatable Prodrugs Based on an Alkoxyamine Platform to Deliver Alkyl Radicals Cytotoxic to Tumor Cells. J Med Chem. 2022;65(13):9253–66. Epub 20220628. doi: 10.1021/acs.jmedchem.2c00455. PubMed PMID: 35764297; PubMed Central PMCID: PMCPMC9289877.

38. Zhang X, Wu L, Zhen W, Li S, Jiang X. Generation of singlet oxygen via iron-dependent lipid peroxidation and its role in Ferroptosis. Fundam Res. 2022;2(1):66–73. Epub 20210809. doi: 10.1016/j.fmre.2021.07.008. PubMed PMID: 38933913; PubMed Central PMCID: PMCPMC11197759.

39. Mercer AE, Maggs JL, Sun XM, Cohen GM, Chadwick J, O’Neill PM, et al. Evidence for the involvement of carbon-centered radicals in the induction of apoptotic cell death by artemisinin compounds. J Biol Chem. 2007;282(13):9372–82. Epub 20070116. doi: 10.1074/jbc.M610375200. PubMed PMID: 17227762.

40. Moore RE, Kim Y, Philpott CC. The mechanism of ferrichrome transport through Arn1p and its metabolism in Saccharomyces cerevisiae. Proc Natl Acad Sci U S A. 2003;100(10):5664–9. Epub 20030429. doi: 10.1073/pnas.1030323100. PubMed PMID: 12721368; PubMed Central PMCID: PMCPMC156258.

41. Philpott CC, Protchenko O. Response to iron deprivation in Saccharomyces cerevisiae. Eukaryot Cell. 2008;7(1):20–7. Epub 20071109. doi: 10.1128/EC.00354-07. PubMed PMID: 17993568; PubMed Central PMCID: PMCPMC2224162.

42. Morgan B, Lahav O. The effect of pH on the kinetics of spontaneous Fe(II) oxidation by O2 in aqueous solution--basic principles and a simple heuristic description. Chemosphere. 2007;68(11):2080–4. Epub 20070321. doi: 10.1016/j.chemosphere.2007.02.015. PubMed PMID: 17368726.

43. Phaniendra A, Jestadi DB, Periyasamy L. Free radicals: properties, sources, targets, and their implication in various diseases. Indian J Clin Biochem. 2015;30(1):11–26. Epub 20140715. doi: 10.1007/s12291-014-0446-0. PubMed PMID: 25646037; PubMed Central PMCID: PMCPMC4310837.

44. Valko M, Leibfritz D, Moncol J, Cronin MT, Mazur M, Telser J. Free radicals and antioxidants in normal physiological functions and human disease. Int J Biochem Cell Biol. 2007;39(1):44–84. Epub 20060804. doi: 10.1016/j.biocel.2006.07.001. PubMed PMID: 16978905.

45. Murphy MP. How mitochondria produce reactive oxygen species. Biochem J. 2009;417(1):1–13. doi: 10.1042/BJ20081386. PubMed PMID: 19061483; PubMed Central PMCID: PMCPMC2605959.

46. Redza-Dutordoir M, Averill-Bates DA. Activation of apoptosis signalling pathways by reactive oxygen species. Biochim Biophys Acta. 2016;1863(12):2977–92. Epub 20160917. doi: 10.1016/j.bbamcr.2016.09.012. PubMed PMID: 27646922.

47. Carneiro BA, El-Deiry WS. Targeting apoptosis in cancer therapy. Nature Reviews Clinical Oncology. 2020;17(7):395–417. doi: 10.1038/s41571-020-0341-y.

48. Bourez S, Van den Daelen C, Le Lay S, Poupaert J, Larondelle Y, Thomé J-P, et al. The dynamics of accumulation of PCBs in cultured adipocytes vary with the cell lipid content and the lipophilicity of the congener. Toxicology Letters. 2013;216(1):40–6. doi: 10.1016/j.toxlet.2012.09.027.

49. Mundy WR, Freudenrich TM, Crofton KM, DeVito MJ. Accumulation of PBDE-47 in Primary Cultures of Rat Neocortical Cells. Toxicological Sciences. 2004;82(1):164–9. doi: 10.1093/toxsci/kfh239.

50. Stadnicka-Michalak J, Tanneberger K, Schirmer K, Ashauer R. Measured and modeled toxicokinetics in cultured fish cells and application to in vitro-in vivo toxicity extrapolation. PLoS One. 2014;9(3):e92303. Epub 20140319. doi: 10.1371/journal.pone.0092303. PubMed PMID: 24647349; PubMed Central PMCID: PMCPMC3960223.

51. Schreiber R, Altenburger R, Paschke A, Küster E. How to deal with lipophilic and volatile organic substances in microtiter plate assays. Environmental Toxicology and Chemistry. 2008;27(8):1676–82. doi: 10.1897/07-504.1.

52. Fischer FC, Cirpka OA, Goss KU, Henneberger L, Escher BI. Application of Experimental Polystyrene Partition Constants and Diffusion Coefficients to Predict the Sorption of Neutral Organic Chemicals to Multiwell Plates in in Vivo and in Vitro Bioassays. Environ Sci Technol. 2018;52(22):13511–22. Epub 20181031. doi: 10.1021/acs.est.8b04246. PubMed PMID: 30298728.

53. Faravelli T, Pinciroli M, Pisano F, Bozzano G, Dente M, Ranzi E. Thermal degradation of polystyrene. Journal of Analytical and Applied Pyrolysis. 2001;60(1):103–21. doi: 10.1016/S0165-2370(00)00159-5.

54. Fujii H, Kakinuma K. Direct Measurement of Superoxide Anion Produced in Biological Systems by ESR Spectrometry: A pH-Jump Method. The Journal of Biochemistry. 1990;108(6):983–7.

